# Itaconic acid production from acetate by *Ustilago maydis*: A step towards land-free biotechnology

**DOI:** 10.64898/2026.01.30.702788

**Authors:** Andreas Müsgens, Lina Wilke, Lars M. Blank

## Abstract

Itaconic acid is a versatile bio-based platform chemical produced from sugar-based feedstocks, linking its production to arable land use. As global food demand rises, alternative carbon sources that decouple industrial biotechnology from agriculture are required. The C_2_ compound acetate can be derived from lignocellulosic biomass and industrial side streams. Emerging routes enable the direct synthesis of acetate from C_1_ carbon sources such as CO_2_, CO, and methane. Here, we show that the smut fungus *Ustilago maydis* can efficiently produce itaconic acid using acetate as the sole carbon source. To overcome weak-acid toxicity and pH-related stress, a combined pH-stat and DO-triggered feeding strategy was applied in a 1 L-scale fed-batch bioreactor, enabling an itaconate titer of 97 g L^−1^ and an overall yield of 0.41 g g^−1^. Key performance indicators were comparable to those of a glucose-based reference process. Despite substantially lower biomass formation on acetate, biomass-specific production rates were markedly higher than on glucose, indicating highly efficient channeling of carbon toward product formation. Overall, our results establish acetate as a competitive and sustainable feedstock for fungal itaconic acid production and position acetate-based processes as a viable route toward land-free biotechnology.

## 1. Introduction

Itaconic acid (ITA) is an unsaturated C_5_ dicarboxylic acid. Due to its structure, it serves as a flexible bio-based platform for the chemical industry. The presence of a vinyl group, along with two carboxyl groups, allows ITA to function as a monomer or comonomer in various polymer applications (Devi et al., 2022). Itaconic acid can replace petrochemical building blocks, such as acrylic and methacrylic acids, in resins, dispersions, superabsorbents, fibers, and coatings (Kuenz and Krull, 2018; Steiger et al., 2017; Teleky and Vodnar, 2019; Werpy and Petersen, 2004). As demand for bio-based polymers and functional materials rises, there is growing industrial interest in robust, cost-effective microbial processes for the production of ITA.

Today, industrial ITA production relies on aerobic submerged fermentation with the filamentous fungus *Aspergillus terreus* on sugar-based substrates (Willke and Vorlop, 2001). These fermentations reach titers of around 160 g L^−1^ on glucose in fed-batch fermentations. However, the bioprocess with *Aspergillus* has its challenges (Krull et al., 2017). Filamentous growth increases fermentation broth viscosity, reducing oxygen transfer, while the mycelium is sensitive to shear forces, forcing operation within a narrow window that balances oxygen supply and mechanical stress (Klement et al., 2012). To overcome these inherent limitations of filamentous morphology, smut fungi of the genus *Ustilago*, which are like Aspergillus native itaconate producers, have emerged as attractive hosts for ITA production. The major advantage of *Ustilago* is its ability to grow in a yeast-like, unicellular state, simplifying process control and scale-up. In *U. maydis*, deletion of the MAP kinase gene *fuz7* abolishes filamentous growth even under challenging growth conditions, leading to a stable yeast-like morphology (Tehrani, 2019).

At the metabolic level, in both *Aspergillus* and *Ustilago*, itaconate is derived from the tricarboxylic acid (TCA) cycle intermediate cis-aconitate, which is exported from mitochondria into the cytosol by dedicated transporters (Agrimi et al.). In *A. terreus*, cis-aconitate is directly decarboxylated to itaconate by the cis-aconitate decarboxylase (CadA). *U. maydis* uses a two-step route in which aconitate-Δ-isomerase (Adi1) first converts cis-aconitate into trans-aconitate, followed by decarboxylation through the trans-aconitate decarboxylase (Tad1) (Geiser et al., 2016b; Wierckx et al., 2020). Over the past years, *U. maydis* has been engineered into a capable chassis for itaconate production. Deletion of competing pathways, combined with overexpression of the cluster regulator Ria1, markedly increased itaconate formation, while additional overexpression of the *A. terreus* cis-aconitate transporter MttA in *U. maydis* further boosted production (Geiser et al., 2016a; Tehrani et al., 2019b). At the end of this development, current *U. maydis* chassis strains operate near the theoretical maximum yield (Becker et al., 2020a; Becker et al., 2020b). In situ product precipitation allows titers of up to 220 g L^−1^ (Tehrani et al., 2019a).

The choice of carbon source is a central lever for improving sustainability and reducing the costs of ITA production. To support a sustainable bioeconomy, microbial processes should be decoupled from arable land and food-grade biomass by relying on non-food, waste-derived, or C_1_/C_2_ feedstocks (Blank et al., 2020). Acetate fits this concept. It is released during lignocellulosic biomass pretreatment, is a major component of lignocellulosic hydrolysates and biorefinery side streams (Kiefer et al., 2021). In addition, acetate can be produced directly from CO_2_ via gas fermentation with acetogenic bacteria (Kantzow et al., 2015), by green chemistry through the isomerization of methyl formate (Jürling-Will et al., 2022), and in bioelectrochemical systems (Rabaey and Rozendal, 2010). Acetate is positioned at the interface of power-to-X chemistry and microbial production (Novak and Pflügl, 2018; Venkata Mohan et al., 2016). These routes offer a genuinely “land-free” and potentially low-cost acetate supply (Mauri et al., 2025). Beyond these emerging sustainable sources, acetate is already available from petrochemical processes (e.g., methanol carbonylation, oxidation of light hydrocarbons) and established fermentations, such as vinegar production on a sugar of biomass basis (Hutkins, 2006). Although these conventional routes are less aligned with the “land-free” vision, they underline that acetate is a mature, large-volume platform molecule. Taken together, the combination of fossil, biomass-derived, and CO_2_-based routes positions acetate as a versatile carbon mediator that can bridge current fossil-dependent processes and future renewable, land-free feedstocks for industrial biotechnology.

Techno-economic analyses of ITA production processes identified substrate costs and product yield as the major contributors to operating expenses (Ernst et al., 2024a; Saur et al., 2023). Recent commodity price data suggest that sodium acetate and acetic acid are typically traded in the range of 0.3-0.6 US$ kg^−1^, while refined sugar prices are on the order of ∼0.35 US$ kg^−1^, indicating a comparable price level between C_2_- and sugar-based feedstocks (Business Analytiq; Trading Economics, 2026). Even if alternative substrates reduce costs for pH-control reagents or wastewater treatment, such savings cannot compensate for substantial losses in carbon conversion efficiency. Consequently, low-cost substrates such as acetate are only attractive if high product yields and space–time yields can be maintained at levels comparable to glucose-based benchmarks.

Despite its promise as a sustainable carbon source, acetate utilization poses known challenges from both a physiological and process-engineering perspective. As a weak acid, acetate can cross biological membranes in its protonated form and dissociate in the cytosol, leading to intracellular acidification and increased maintenance demands (Kutscha and Pflügl, 2020; Romero-Aguilar et al., 2023). In addition, during itaconate production from acetate, the pH increases because the overall reaction, for simplicity assuming 100% yield, converts three acetate ions into one itaconate dianion and CO_2_, requiring net proton uptake from the medium. The CO_2_ is stripped from the liquid phase in an aerated system, further intensifying this trend.

Recent work by Merkel et al. (2022) demonstrated that these constraints can be addressed by a tailored dual fed-batch strategy for itaconate production from acetate in *Corynebacterium glutamicum*. In their process, glacial acetic acid served as both a pH titrant and a carbon feed. Merkel et al. implemented a second DO-controlled feed of sodium acetate to replenish acetate in the event of carbon depletion. A purely pH-stat-based strategy would be insufficient because the product is an acid, so titrant addition will always run towards zero.

In this study, we evaluated acetate as the sole carbon source for itaconic acid production by *Ustilago maydis*. We first characterized acetate utilization and production performance of the latest *U. maydis* itaconate chassis strain in small-scale batch cultivation to capture physiological constraints and pH dynamics. We then scaled the process to 0.8 L bioreactors operated in fed-batch mode, following the conceptual process blueprint of Merkel et al. (2022). Finally, we benchmarked the acetate process against a glucose-fed reference fermentation. Overall, this work aims to establish *U. maydis* as an efficient itaconate producer on acetate and to further advance acetate as a sustainable C_2_ feedstock for industrial biotechnology.

## 2. Materials and Methods

### 2.1. Growth media and cultivation conditions

YPD-agar was used to cultivate *U. maydis* from glycerol stocks for the inoculation of precultures. The medium contained (per L) 20 g agar, 10 g peptone, 20 g glucose, 10 g yeast extract. Agar plates were grown at 30 °C for two days.

Itaconate production experiments were performed in minimal Modified Tabuchi Medium (MTM) (Geiser et al., 2016a) containing (per L) 0.2 g MgSO_4_·7 H_2_O, 0.01 g FeSO_4_·7 H_2_O, 0.5 g KH_2_PO_4_, 1 mL trace element solution, 1 mL vitamin solution, and as buffer either 3-(N-morpholino)propanesulfonic acid (MOPS) or 2-(N-morpholino)ethanesulfonic acid (MES) at the concentrations specified for each experiment. Carbon sources (glucose, acetate) and the nitrogen source (NH_4_Cl) varied according to the experimental setup and are reported in the respective sections. The vitamin solution contained (per L) 0.05 g D-biotin, 1 g D-calcium pantothenate, 1 g nicotinic acid, 25 g myo-inositol, 1 g thiamine hydrochloride, 1 g pyridoxal hydrochloride, and 0.2 g para-aminobenzoic acid. The trace element solution contained (per L) 1.5 g EDTA, 0.45 g ZnSO_4_ 7H_2_O, 0.10 g MnCl_2_ 4H_2_O, 0.03 g CoCl_2_ 6H_2_O, 0.03 g CuSO4·5H_2_O, 0.04 g Na_2_MoO_4_ 2H_2_O, 0.45 g CaCl_2_ 2H_2_O, 0.3 g FeSO_4_ 7H_2_O, 0.10 g H_3_BO_3_, and 0.01 g KI.

### 2.2. Shake flask cultivation

Itaconate production strain *U. maydis Δfuz7 Δcyp3 ΔMEL ΔUA P*_*ria1*_*::P*_*etef*_ *P*_*etef*_*mttA* (iAMB collection #7078; Ullmann, 2023) was characterized for growth and production on acetate. Precultures were inoculated from agar plates and grown in MTM containing 50 g L^-1^ glucose, 2 g L^-1^ acetate (as sodium acetate), 1.6 g L^-1^ NH_4_Cl, and 19.5 g L^-1^ MES at a starting pH of 6.5. Precultures were cultivated at 30 °C with 15 mL filling volume in 100 mL flasks for 24 h. Cell material was washed twice before inoculation of the main culture by pelleting at 4000 g for 5 min and resuspending the pellets in 0.9% NaCl.

The main culture was inoculated with an initial OD_600_ of 0.5 in MTM containing 4 g L^-1^ acetate (sodium salt), 0.1 g L^-1^ NH_4_Cl, and 10.47 g L^-1^ 3-(N-morpholino)propanesulfonic acid (MOPS), adjusted to pH 7.5. Cultivations were performed at 30 °C in a 500 mL shake flask with 50 mL filling volume, shaken at 200 rpm with 25 mm throw.

### 2.3. Fed-batch bioreactor cultivation on acetate

Fed-batch cultivations on acetate were performed in a BioFlo® 120 2 L bioreactor with a starting volume of 0.8 L controlled via DASware Control Software (Eppendorf, Hamburg, Germany). Cultivations were carried out in MTM medium (3 g L^-1^ acetate, 4 g L^-1^ NH_4_Cl, 10.46 g L^-1^ MOPS) with an initial pH of 7.0. The pH was monitored with an online probe (EasyFerm Plus PHI K8 255, Hamilton, Bonaduz, Switzerland) and maintained at pH 6.8 by titration with glacial acetic acid, which also served as carbon feed. Dissolved oxygen (DO) was monitored using an InPro 6850 electrode (Mettler Toledo, Columbus, USA) and maintained at 70 to 80% air saturation by manual adjustment of the agitation speed (400-1,200 rpm). A DO-based automated script (Supplementary Note S1) triggered the addition of sodium acetate (200 g acetate per L) solution in case of carbon depletion. Each feed pulse restored the acetate concentration to 4 g L^-1^. The bioreactor was aerated at 2-3 L min^-1^, and evaporation was minimized by sparging the inlet gas through a water bottle. Off-gas CO_2_ and O_2_ concentrations were measured using a BlueVary Sensor and BlueVis software (BlueSens gas sensor GmbH, Herten, Germany). The temperature was set at 30 °C. The bioreactor was inoculated to a final OD_600_ of 1 with cells from an overnight culture in 50 mL MTM containing 50 g L^-1^ glucose, 2 g L^-1^ acetate, 1.6 g L^-1^ NH_4_Cl, and 19.5 g L^-1^ MES (pH 6.5). Antifoam was added manually when required. The weight of the feed/pH-titrant flasks was recorded for each sampling point. The process design was conceptually adapted from Merkel et al. (2022).

### 2.4. Fed-batch bioreactor cultivation on glucose

Fed-batch cultivations on glucose were performed in a BioFlo® 120 2 L bioreactor with a starting volume of 0.8 L controlled via DASware Control Software (Eppendorf, Hamburg, Germany). Cultivations were carried out in MTM medium (10 g L^-1^ glucose at the start, 4 g L^-1^ NH_4_Cl, 9.76 g L^-1^ MES) with an initial pH of 6.5. The pH was monitored with an online probe (EasyFerm Plus PHI K8 255, Hamilton, Bonaduz, Switzerland) and maintained at pH 6.5 by automatic addition of 10 M NaOH. Dissolved oxygen (DO) was monitored using an InPro 6850 electrode (Mettler Toledo, Columbus, USA) and maintained at 70 to 80% air saturation by manual adjustment of the agitation speed (400-1,200 rpm). A DO-based automated script (Supplementary Note S1) triggered the addition of a glucose (500 g L^-1^) solution upon carbon depletion. Each feed pulse restored the glucose concentration to 15 g L^-1^. The bioreactor was aerated at 2 L min^-1^, and evaporation was minimized by sparging the inlet gas through a water bottle. Off-gas CO_2_ and O_2_ concentrations were measured using a BlueVary Sensor and BlueVis software (BlueSens gas sensor GmbH, Herten, Germany). The temperature was set at 30 °C. The bioreactor was inoculated to a final OD_600_ of 1 with cells from an overnight culture in 50 mL MTM containing 50 g L^-1^ glucose, 1.6 g L^-1^ NH_4_Cl, and 19.5 g L^-1^ MES (pH 6.5). Antifoam was added manually when required. The weight of the feed and pH-titrant flasks was recorded for each sampling point.

### 2.5. Metabolite quantification via HPLC-UV/RI

Acetate, glucose, and itaconate were quantified using an Ultimate 3000 HPLC system (Thermo Scientific, Waltham, USA). Samples were centrifuged (2 min, 14,000 rpm), and the supernatants were stored at -20°C for HPLC measurement. Supernatants were diluted prior to measurement (typically 2- to 10-fold). All samples were filtered through 0.2 µm syringe filters. Separation was achieved under isocratic conditions with 5 mM H_2_SO_4_ as the mobile phase at a flow rate of 0.6 mL min^-1^. A 5 µL injection volume was applied onto a Metab-AAC ion-exchange column (300 × 7.8 mm, 10 µm, Isera GmbH, Düren, Germany) maintained at 40 °C. Metabolites were quantified using UV (210 nm) and refractive index detectors.

### 2.6. Optical density measurement

Cell growth was monitored by measuring the optical density at 600 nm with an Ultrospec 10-cell Density Meter (Amersham Biosciences; Amersham, UK).

### 2.7. Cell dry weight determination

Cell dry weight (CDW) was determined in triplicate. For CDW determination, 2 mL of the fermentation broth were transferred into a pre-dried, pre-weighed tube and centrifuged at 12,000 rpm for 5 min. The cell pellet was washed by resuspension in 1 mL of 0.9% (w/v) NaCl. After a second centrifugation step (5 min, 12,000 rpm), the supernatant was removed, and the cell pellet was dried in a vacuum oven for 72 h at 60 °C. The CDW was determined as the weight difference between the empty tube and the tube containing the dried pellet.

### 2.8. Statistical analysis

For the characterization of itaconate production in shake-flask batch cultivations (Section 3.1), the reported values represent the mean of a biological triplicate. Bioreactor cultivations were performed in duplicate. For duplicate experiments, the reported values correspond to the mean, with the given error indicating the minimum and maximum values.

## 3. Results

### 3.1 Small-scale characterization of acetate utilization and production strain performance

The itaconate production strain *U. maydis Δfuz7 Δcyp3 ΔMEL ΔUA P*_*ria1*_*::P*_*etef*_ *P*_*etef*_*mttA* was evaluated regarding growth and production performance on acetate in a small-scale batch cultivation (Fig. 1). A low initial acetate concentration of 4 g L^-1^ was chosen, combined with 0.1 g L^-1^ NH_4_Cl, to trigger nitrogen limitation and initiate itaconate production even with limited C-source availability. To reduce weak-acid stress, the initial pH was elevated from 6.5 to 7.5 compared to standard conditions on glucose. Detailed sampling of the shake flask showed that the pH rose from 7.5 to 9.3 within 48 h, coincident with the itaconate production phase, revealing a rapid pH increase despite an already optimized buffer system. Acetate was consumed after approximately 66 h, ending itaconate production. The final itaconate titer reached 1.66 ± 0.4 g L^-1^ with a yield of 0.42 ± 0.01 g g^-1^. These results demonstrate that the strain efficiently converts acetate even under the inherent limitations of a batch culture. To the best of our knowledge, itaconate production with *Ustilago maydis* from acetate as the sole carbon source has not been previously described.

**Figure 1.**
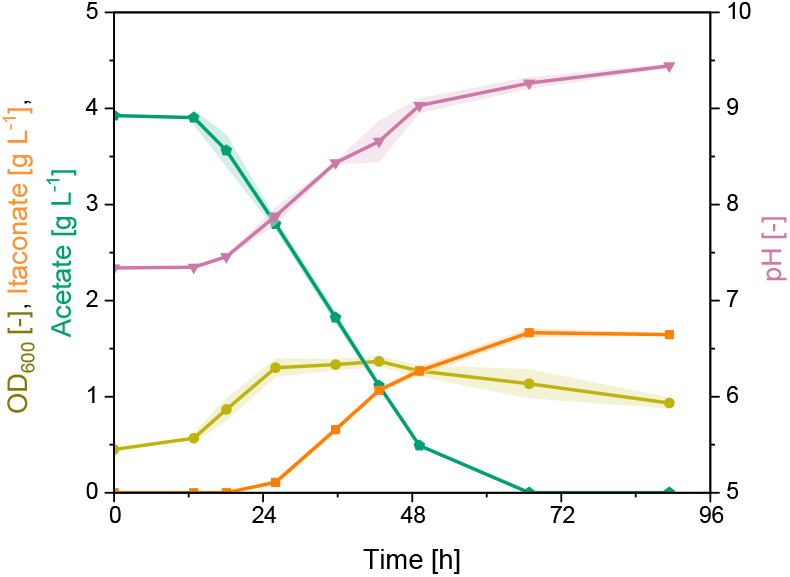
Shake flask cultivation of *Ustilago maydis* on acetate. Itaconate concentration (orange squares), optical density (OD) (green circles), acetate concentration (teal pentagons), and pH (violet triangles) are plotted over cultivation time. The cultivation was performed at 30°C in MTM medium (50 mM MOPS, pH 7.5, 0.1 g L^-1^ NH_4_Cl). Shaking conditions: 50 mL in a 500 mL flask, 200 rpm, 25 mm throw. Itaconate production strain *U. maydis Δfuz7 Δcyp3 Δ*MEL *Δ*UA P_ria1_::P_etef_ P_etef_mttA was used as the production host. Shades represent the standard deviation of a biological triplicate (n=3).

### 3.2 Fed-batch bioreactors with acetate as carbon source

Shake flask experiments revealed three main bottlenecks for itaconate production from acetate in batch mode: (i) slow growth, (ii) acetate-induced inhibition through weak-acid toxicity, and (iii) a rapid pH increase despite high buffer capacity. In a bioreactor, these limitations can be overcome through pH control and a fed-batch strategy.

Consequently, we performed a fed-batch bioreactor cultivation in duplicate (Fig. 2), starting with a cultivation volume of 0.8 L in minimal medium. Targets for process control were to maintain acetate concentrations below 4 g L^-1^ and to keep the pH constant at 6.8, slightly higher than with our reference substrate, glucose, to reduce weak acid toxicity during the production phase. A dual-feed strategy was implemented, with glacial acetic acid serving as both pH titrant and feed solution, while sodium acetate additionally supplied the carbon source via DO-triggered pulses in case of carbon depletion. The time point of nitrogen source depletion can be pinpointed to 33 h after cultivation start, as indicated by off-gas data (not shown), marking the end of the growth phase and the beginning of the production phase. Cell dry weight (CDW) stabilized around 13 g L^-1^ (Supplemental Fig. 1) during the production phase. Over the 262 h total cultivation time, 456 g acetate were consumed, 268 g as acetic acid, and 188 g in the form of sodium acetate. Total itaconate production reached 188 ± 19 g, corresponding to a titer of 96.8 ± 7.7 g L^-1^. The overall yield achieved was 0.41 ± 0.03 g g^-1^, and the space-time yield averaged 0.39 ± 0.03 g L^-1^ h^-1^, with a peak productivity of 1.04 ± 0.05 g L^-1^ h^-1^ between 44 h and 70 h. The reactor volume increased from 0.79 L to 1.83 L during cultivation. After 170 h, itaconate production declined, likely due to the additive effects of Na^+^ accumulation through the sodium acetate feed and product inhibition. A summary of the key process parameters is given in Table 1. To our knowledge, the data shown here represent the first successful production of itaconate by *Ustilago maydis* using acetate as the sole carbon source in a controlled bioreactor setup.

**Table 1.**
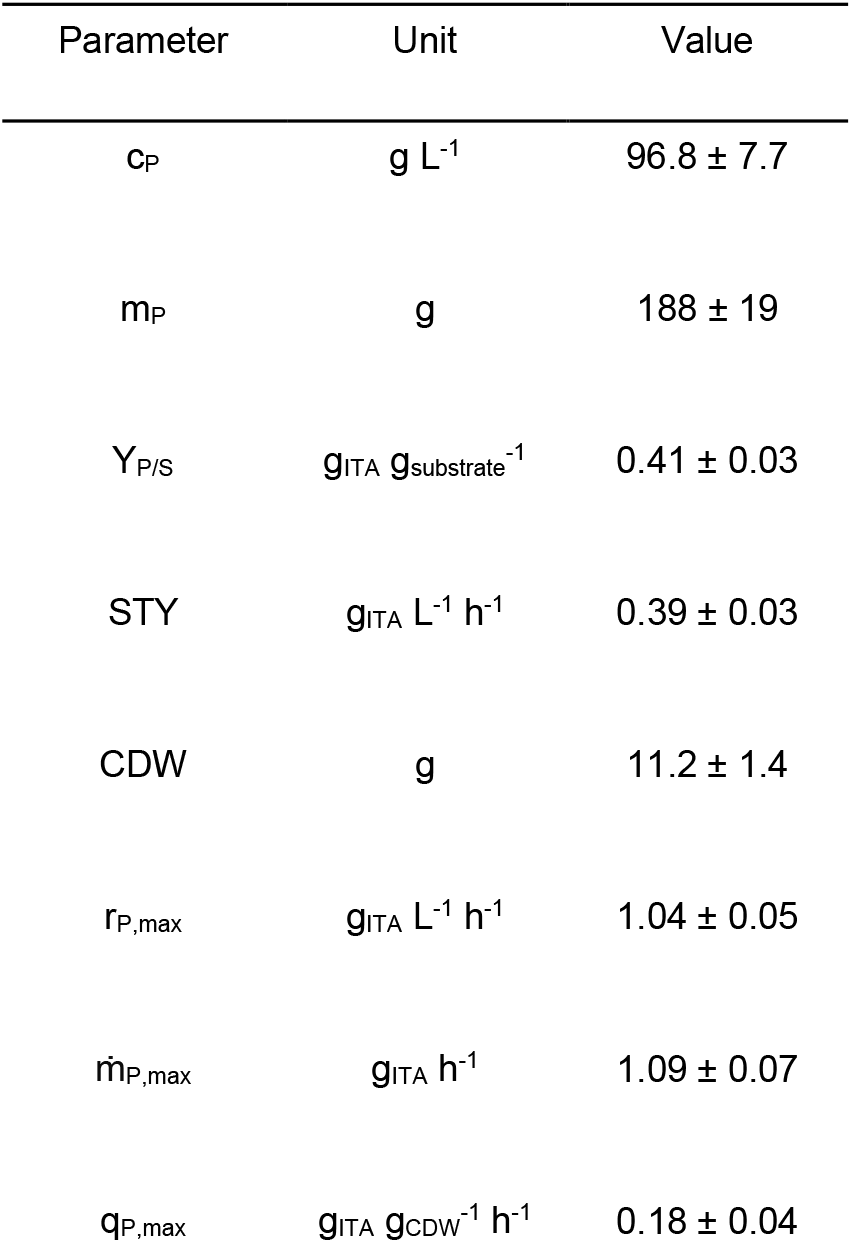
Overview of key performance parameters for the fed-batch cultivation on acetate (n=2). With c_P_: final product concentration, m_P_: final product mass, Y_P/S_: final product-to-substrate yield based on final product mass and consumed substrate, STY: overall space-time yield calculated from final product concentration divided by the total process time, CDW: cell dry weight represents the total biomass formed, r_P,max_: maximum volumetric production rate between two sampling points, 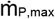: maximum absolute production rate between two sampling points, and q_P,max_: maximum biomass-specific production rate between two sampling points. Errors represent the minimum and maximum values of a biological duplicate

| Parameter | Unit | Value |
| --- | --- | --- |
| $c_P$ | $g\ L^{-1}$ | $96.8 \pm 7.7$ |
| $m_P$ | g | $188 \pm 19$ |
| $Y_{P/S}$ | $g_{ITA}\ g_{substrate}^{-1}$ | $0.41 \pm 0.03$ |
| STY | $g_{ITA}\ L^{-1}\ h^{-1}$ | $0.39 \pm 0.03$ |
| CDW | g | $11.2 \pm 1.4$ |
| $r_{P,max}$ | $g_{ITA}\ L^{-1}\ h^{-1}$ | $1.04 \pm 0.05$ |
| $\dot{m}_{P,max}$ | $g_{ITA}\ h^{-1}$ | $1.09 \pm 0.07$ |
| $q_{P,max}$ | $g_{ITA}\ g_{CDW}^{-1}\ h^{-1}$ | $0.18 \pm 0.04$ |

**Figure 2.**
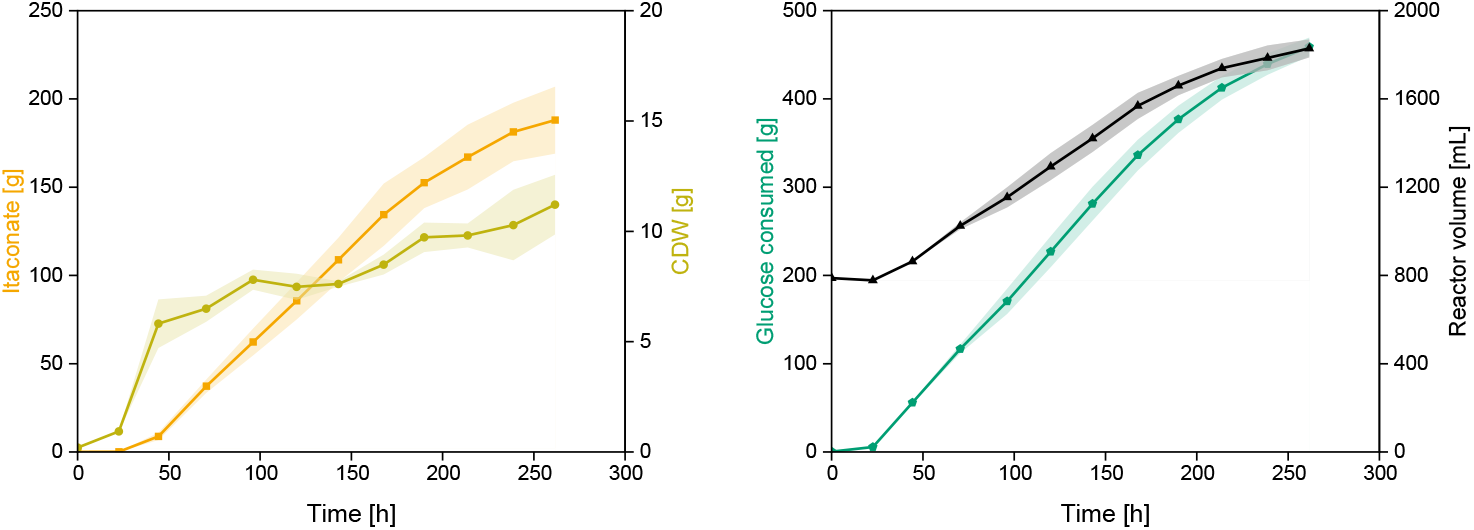
Bioreactor cultivation of *Ustilago maydis* for itaconate production on acetate using a pH and DO-coupled feeding strategy. Itaconate (orange squares), cell dry weight (CDW) (green circles), acetate consumption (teal pentagons), and reactor volume (black triangles) are plotted over the cultivation time. The cultivation was performed at 30°C in MTM medium (50 mM MOPS, pH 6.8, 4 g L^-1^ NH_4_Cl) with dissolved oxygen levels between 70 and 80%. The pH was controlled with glacial acetic acid as titrant. DO-coupled fed shots with sodium acetate (200 g_ACE_ L^-1^), replenished the glucose concentration to 4 g L^-1^ in case of carbon depletion. Itaconate production strain *U. maydis Δfuz7 Δcyp3 Δ*MEL *Δ*UA P_ria1_::P_etef_ P_etef_mttA was used as the production host. Shades represent the minimum and maximum values of a biological duplicate (n=2).

### 3.3 Glucose reference fermentation

For benchmarking, a fed-batch process on glucose was carried out in duplicate (Fig. 3, Table 2), mirroring the acetate cultivation as closely as possible. On glucose, the pH does not increase during the production phase but instead shifts towards lower values. Consequently, the pH titrant was changed from glacial acetic acid to 10 M NaOH. The pH setpoint during the production phase was adjusted to the standard production value of 6.5. Feeding was controlled exclusively via DO-triggered pulses using a 500 g L^-1^ glucose solution as the reservoir, restoring the glucose concentration in the cultivation medium to about 15 g L^-1^ with each feeding event.

**Table 2.** Overview of key performance parameters for the fed-batch cultivation on glucose (n=2). With c_P_: final product concentration, m_P_: final product mass, Y_P/S_: final product-to-substrate yield based on final product mass and consumed substrate, STY: overall space-time yield calculated from final product concentration divided by the total process time, CDW: cell dry weight represents the total biomass formed, r_P,max_: maximum volumetric production rate between two sampling points, 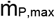 : maximum absolute production rate between two sampling points, and q_P,max_: maximum biomass-specific production rate between two sampling points. Errors represent the minimum and maximum values of a biological duplicate

| Parameter | Unit | Value |
| --- | --- | --- |
| $c_P$ | $g\ L^{-1}$ | $80.8 \pm 2.4$ |
| $m_P$ | $g$ | $117.7 \pm 3.6$ |
| $Y_{P/S}$ | $g_{ITA}\ g_S^{-1}$ | $0.41 \pm 0.01$ |
| STY | $g_{ITA}\ L^{-1}\ h^{-1}$ | $0.46 \pm 0.01$ |
| CDW | $g$ | $19.1 \pm 0.3$ |
| $r_{P,max}$ | $g_{ITA}\ L^{-1}\ h^{-1}$ | $1.04 \pm 0.02$ |
| $\dot{m}_{P,max}$ | $g_{ITA}\ h^{-1}$ | $1.08 \pm 0.02$ |
| $q_{P,max}$ | $g_{ITA}\ g_{CDW}^{-1}\ h^{-1}$ | $0.11 \pm 0.01$ |

**Figure 3.**
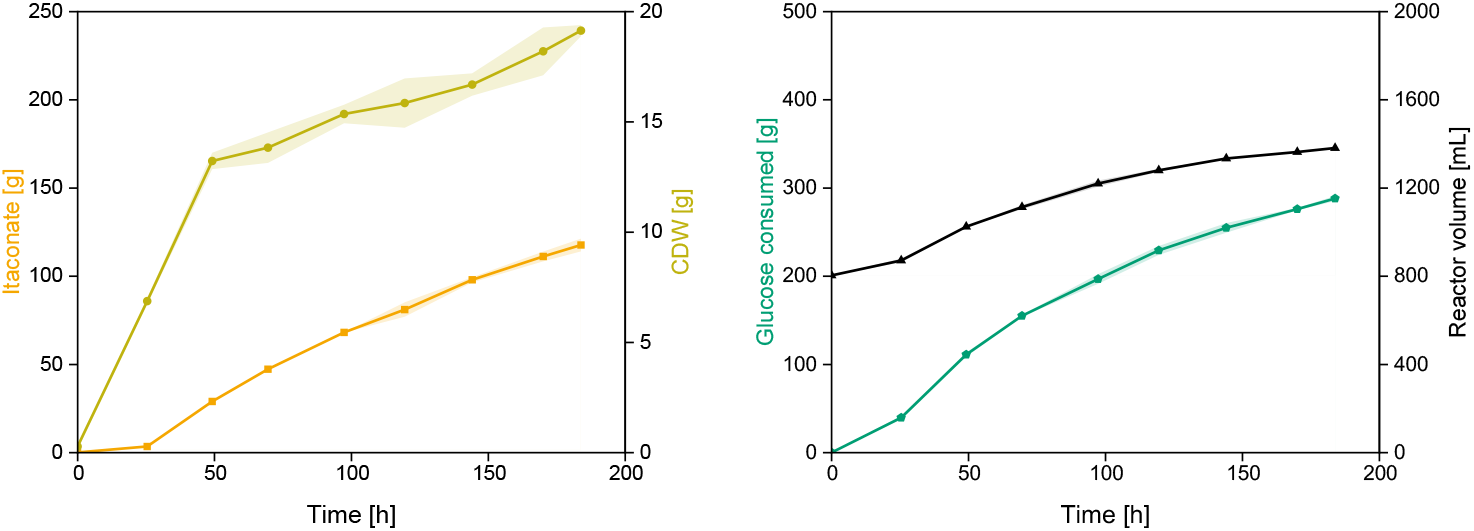
Bioreactor cultivation of *Ustilago maydis* for itaconate production on glucose using a DO-coupled feeding strategy. Itaconate (orange squares), cell dry weight (CDW) (green circles), glucose consumption (teal pentagons), and reactor volume (black triangles) are plotted over the cultivation time. The cultivation was performed at 30°C in MTM medium (50 mM MES, pH 6.5, 4 g L^-1^ NH_4_Cl) with dissolved oxygen levels between 70 and 80%. The pH was controlled with 10 M NaOH. DO-coupled fed with glucose (500 g L^-1^), replenished the glucose concentration back to 15 g L^-1^ in case of carbon depletion. Itaconate production strain *U. maydis Δfuz7 Δcyp3 Δ*MEL *Δ*UA P_ria1_::P_etef_ P_etef_mttA was used as the production host. Shades represent the minimum and maximum values of a biological duplicate (n=2).

According to off-gas data (not shown), nitrogen was depleted after 22 h. Biomass concentration stabilized at 12 g L^-1^ and remained constant throughout the run. Over the total process time of 184 h, 117.7 ± 3.6 g itaconate were produced, and 288 g glucose were consumed. A final titer of 80.8 ± 2.4 g L^-1^ was achieved. The overall yield was 0.41 ± 0.01 g g^-1^, and the space-time yield reached 0.46 ± 0.01 g L^-1^ h^-1^, with a maximum volumetric production rate of 1.04 ± 0.02 g L^-1^ h^-1^ between 25 h and 50 h. Reactor volume increased from 0.80 L to 1.38 L during the cultivation.

For a direct comparison of both processes, the time point at 170 h of cultivation was evaluated, representing a reasonable endpoint for both processes due to declining production rates. Key performance indicators after 170 h are listed in Table 3. For the acetate process, 134.4 ± 17.7 g itaconate were produced after 170 h, compared to 111.2 ± 2.8 g on glucose (Fig. 4 A). Product titers were similar, with 82.4 ± 7.8 g L^−1^ for acetate and 78.0 ± 2.0 g L^−1^ for glucose. Overall yields were identical at 0.40 ± 0.03 g g^-1^ for acetate and 0.40 ± 0.01 g g^−1^ for glucose (Fig. 4 B), reflecting comparable carbon-conversion efficiency. Biomass formation was faster on glucose and reaches a higher level despite the same amount of available nitrogen source. After 170 h, the acetate-fed process reached 8.5 ± 0.4 g CDW, while the glucose-fed process accumulated more than twice as much (18.2 ± 1.1 g CDW). Lower biomass formation led to significantly higher specific production rates (Fig. 4 C). The non-volumetric production rate 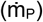 was also slightly higher on acetate throughout the entire production phase (Fig. 4 D).

**Table 3.**
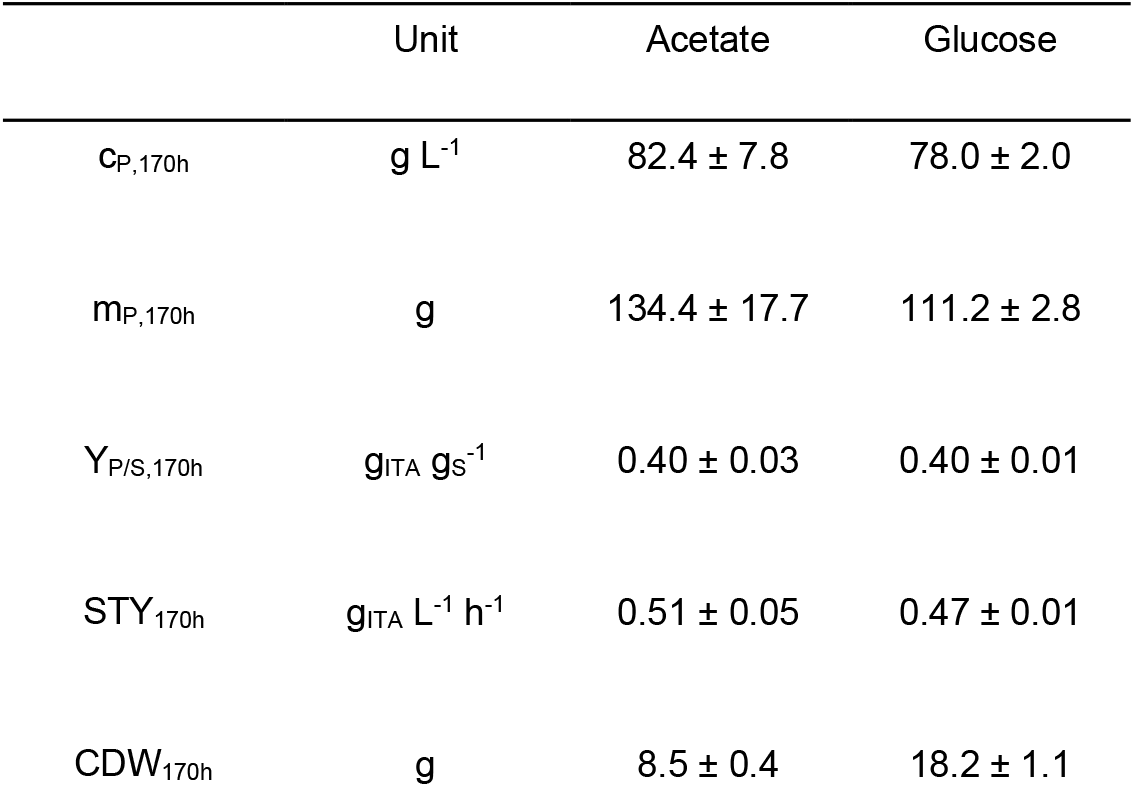
Comparison of key performance parameters for the fed-batch cultivation on acetate and glucose after 170 h of cultivation (n=2). With c_P,170h_: product concentration, m_P,170h_: product mass, Y_P/S,170h_: product-to-substrate yield, STY_170h_: overall space-time yield calculated from product concentration divided by the process time, CDW_170h_: cell dry weight represents the biomass in 170 h. Shades represent the minimum and maximum values of a biological duplicate

**Figure 4.**
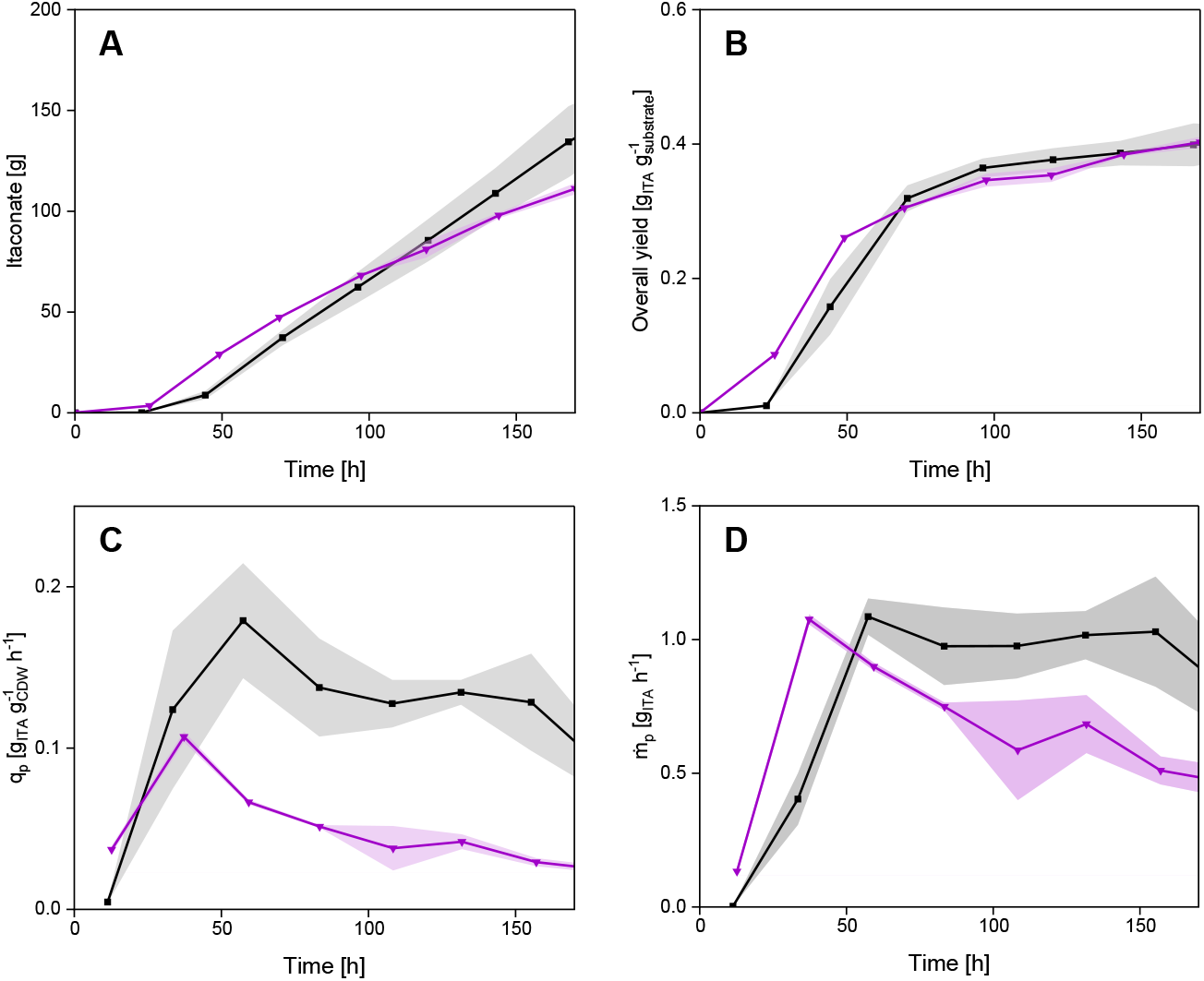
Comparison of process performance for fed-batch bioreactor cultivations on acetate and glucose. Black lines represent the acetate/acid-fed process, and magenta lines the process on glucose. (A) Itaconate formation, (B) overall yield (Y_P/S_), specific production rate (q_P_) and product formation rate 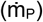 plotted over process time. Shades represent the minimum and maximum values of a biological duplicate (n=2).

These results demonstrate that *Ustilago maydis* maintains a high level of itaconate production on acetate, comparable to that on glucose, across all relevant parameters. Production rates on acetate appear slightly higher, though this may be an artifact of lower stress due to higher dilution in the acetate process. The biomass-specific productivity was substantially higher on acetate, suggesting that achieving biomass concentrations similar to those in the glucose process could further enhance acetate-based production performance.

## 4. Discussion

This study demonstrates that *Ustilago maydis* can produce itaconate from acetate as the sole carbon and energy source. The physiological constraints typically associated with acetate utilization, slow growth, weak-acid stress, and an upward pH shift were clearly observed in shake-flask cultivation. Despite these limitations, the strain efficiently converted acetate into itaconate, achieving a titer of 1.66 g L^−1^ from only 4 g L^−1^ substrate. Transitioning to fed-batch mode mitigated the significant limitations observed in shake flasks by maintaining acetate concentrations at or below 4 g L^−1^ to keep weak-acid toxicity low, while upholding tight pH control. Under these optimized conditions, the acetate-fed process, which was run in duplicate, reached a mean itaconate titer of 96.8 g L^−1^, with one reactor exceeding 100 g L^−1^. The overall yield achieved was 0.41 g g^−1^ with an overall volumetric productivity (STY) of 0.39 g L^−1^ h^−1^. All relevant KPIs after 170 h were at least as high as those in the glucose-fed reference fermentation (Table 3). Notably, the biomass-specific productivity was significantly higher in the acetate-fed fermentation. Given that the maximum stoichiometric yields on glucose and acetate are almost equal (0.72 g g^−1^ and 0.73 g g^−1^, respectively), using a mass basis for yield comparison is appropriate without conversion to C-mol basis. Overall productivity is high, especially considering the long growth phase on acetate.

For comparison, one of the best-reported glucose-based processes with a similar experimental setup using a closely related strain reported an itaconate titer of 76 g L^−1^ with a yield of 0.53 g g^−1^, and a productivity of 0.54 g L^−1^ h^−1^ (Becker et al., 2020a). Itaconate production with *Aspergillus terreus* can achieve 160 g L^−1^, a yield of 0.46 g g^−1^, and a productivity of 0.99 g L^−1^ h^−1^ (Krull et al., 2017), with higher productivity being the main advantage. As in all fed-batch processes, overall volumetric productivity does not benefit from prolonged cultivation targeting high itaconate titers, with ongoing high dilution for the acetate process further adding to the reduced volumetric productivity. The internal glucose reference clearly demonstrates that acetate is not inferior as a substrate for itaconate production and that carbon flux into itaconate is robust.

Compared to conventional second-generation feedstock substrates, acetate performed remarkably well. Itaconate production attempts with *Ustilago cynodontis* on starch did not exceed 10 g L^−1^ in titer and yielded only about 0.1 g g^−1^ (Ernst et al., 2024b). Thick juice, a byproduct of sugar production containing sucrose, was successfully used for itaconate production with *Ustilago*, reaching 106 g L^−1^ in titer with a yield of 0.50 g g^−1^ and a productivity of 0.72 g L^−1^ h^−1^ (Niehoff et al., 2023). Crude glycerol also has potential as a substrate but suffers from low productivity (74.5 g L^−1^, 0.17 g L^−1^ h^−1^; Helm et al., 2024). The high titers, yields, and rates achieved on acetate in the present work therefore position acetate as one of the most promising non-glucose substrates currently evaluated for itaconate production. An overview of further renewable feedstocks and production values can be found in Becker et al. (2022).

Acetate-based itaconate production has thus far been demonstrated only in non-native hosts such as *Escherichia coli* and *Corynebacterium glutamicum. E. coli* achieved 3.6 g L^−1^ in 88 h of cultivation on acetate (Noh et al., 2018), while *C. glutamicum* has shown a high capacity for itaconate production (Merkel et al., 2022). The fed-batch fermentation by Merkel et al. reached an overall titer of 29.2 g L^−1^ with a yield of 0.16 g g^−1^ and a productivity of 0.63 g L^−1^ h^−1^. The peak volumetric productivity of 1.01 g L^−1^ h^−1^ reported in that study is equal to the peak volumetric productivity observed for the acetate process described here. Further strain engineering of *C. glutamicum* led to 3-fold higher yields and 4-fold higher titers in shake-flask cultivations (Schmollack et al., 2022). It is therefore likely that implementing these advanced strains in a fed-batch will further improve itaconate production from acetate with *Corynebacterium*.

A notable physiological difference between the acetate process and glucose reference was the substantially lower biomass formation on acetate, despite identical nitrogen availability. Since cell numbers were not determined, it cannot be excluded that biomass composition differed between processes. Glucose-grown cells may accumulate more energy-dense storage molecules such as neutral lipids, resulting in heavier biomass. Additional metabolic investment required for acetate assimilation, including ATP-dependent substrate activation, gluconeogenesis, and redox balancing, may also be associated with a less efficient use of available nitrogen. The underlying mechanisms remain unclear and require further investigation. The approximately 1.6-fold higher specific production rate observed for the acetate process compared to the glucose reference still offers considerable potential for process optimization. It needs to be investigated whether higher cell densities can be achieved on acetate without causing excessive stress, and whether increased biomass formation in acetate-based processes would indeed translate into enhanced biocatalytic performance.

The acetate fed-batch process experienced substantial dilution due to the limited solubility of sodium acetate (approximately 200 g L^−1^), even though more than half of the fed acetate was supplied in highly concentrated form as 100% glacial acetic acid via the pH control (268 g of 456 g total acetate feed). The culture volume increased from 0.79 to 1.83 L. By comparison, the volume increase for the glucose process was from 0.80 to 1.38 L using a 500 g L^−1^ glucose feed and 10 M NaOH as pH titrant. The resulting dilution had mixed effects: it lowered product titers but also reduced osmotic and weak-acid stress, likely stabilizing production over the long cultivation period. To minimize dilution, a potassium acetate feed was evaluated (Supplemental Fig. 2) due to its higher solubility (>500 g L^−1^). However, despite the more concentrated feed, a potassium acetate-based fed-batch produced only 63 g L^-1^ in 170 h, suggesting that *U. maydi*s is more sensitive to elevated K^+^ than to Na^+^.

In both the glucose and acetate processes, itaconate production began to decline as itaconate concentration increased. This decline due to product inhibition is accompanied by inhibitory effects of salt accumulation and rising osmolarity, which are all frequently observed in organic acid fermentations (Niehoff et al., 2023). Although *Ustilago* species can tolerate high osmotic stress, sodium accumulation is known to impair membrane integrity and stress tolerance in *U. maydis* (Benito et al., 2009; Salmerón-Santiago et al., 2011). Various in situ product removal (ISPR) strategies have demonstrated significant benefits for organic acid fermentations and hold promise for acetate-based itaconate production processes. In situ adsorption using activated carbon resins enables continuous removal of itaconic acid from the fermentation broth, improving overall process performance (Pastoors et al., 2023). Furthermore, membrane-based cell retention and product separation concepts, such as hollow-fiber modules operated in reverse-flow diafiltration mode, enable gentle, selective product recovery at elevated cell densities and have been shown to provide stable operation and enhanced space-time yields in continuous itaconic-acid processes (Carstensen et al., 2013). Fed-batch and continuous setups combining membrane-based cell retention with ISPR represent promising intensification strategies for acetate-driven itaconate bioprocesses. Notably, Tehrani et al. (2019a) reported a record itaconic acid titer of 220 g L^−1^ (Y_P/S_ = 0.33 g g^−1^ and STY = 0.45 g L^−1^ h^−1^) with *U. maydis* by applying an in situ crystallization technique using CaCO_3_ to precipitate itaconic acid during fed-batch fermentation. This strategy effectively reduces product inhibition, enabling extremely high titers.

To address the problem of slow growth on acetate while simultaneously tackling declining productivity in prolonged cultivations, repeated-batch fermentation strategies could be implemented. These approaches allow periodic removal and replacement of the fermentation medium while retaining biomass. Additionally, using glucose as a substrate only during the growth phase should be considered.

The use of acetate offers a clear opportunity to advance towards a more sustainable bioeconomy. Acetate is already abundantly available as a component of lignocellulosic hydrolysates, making it an attractive, low-cost feedstock for industrial biotechnology. Prospective routes for the direct production of acetate from CO_2_ and H_2_ further add to this potential. This study establishes *U. maydis* as an efficient itaconate producer on acetate and shows that this substrate can match glucose in overall process performance. By achieving high titers, robust volumetric productivities, and a carbon-conversion efficiency comparable to glucose, the process presented here positions acetate among the most promising non-glucose substrates for itaconate production and, to the best of our knowledge, reports the highest itaconate titer obtained from acetate with any microorganism to date.

## 5. Author contributions

**Andreas Müsgens**: Conceptualization, Data curation, Investigation, Formal analysis, Methodology, Project administration, Writing – original draft, Writing – review and editing, Visualization

**Lina Wilke**: Methodology, Investigation, Formal analysis.

**Lars M. Blank**: Writing – review & editing, Writing – original draft, Project administration, Supervision, Funding acquisition, Conceptualization

## 6. Acknowledgements

The authors would like to thank their colleagues at the Institute of Applied Microbiology (iAMB) for their continuous and valuable support. This work has received funding from the Deutsche Forschungsgemeinschaft (DFG, German Research Foundation) – SFB1535 - Project ID 458090666. The laboratory of LMB is partially supported by the Deutsche Forschungsgemeinschaft (DFG, German Research Foundation) under Germany’s Excellence Strategy, Cluster of Excellence 2186 “The Fuel Science Center” – ID: 390919832 and by the Werner Siemens Foundation in the frame of the WSS Research Centre “catalaix”.

## 8. Data availability

Data supporting the findings of this work are available from the corresponding author on request.

