## Supplemental Material for "Itaconic acid production from acetate by *Ustilago maydis*: A step towards land-free biotechnology" (from another version of this preprint: version 1, 2026-01-30)

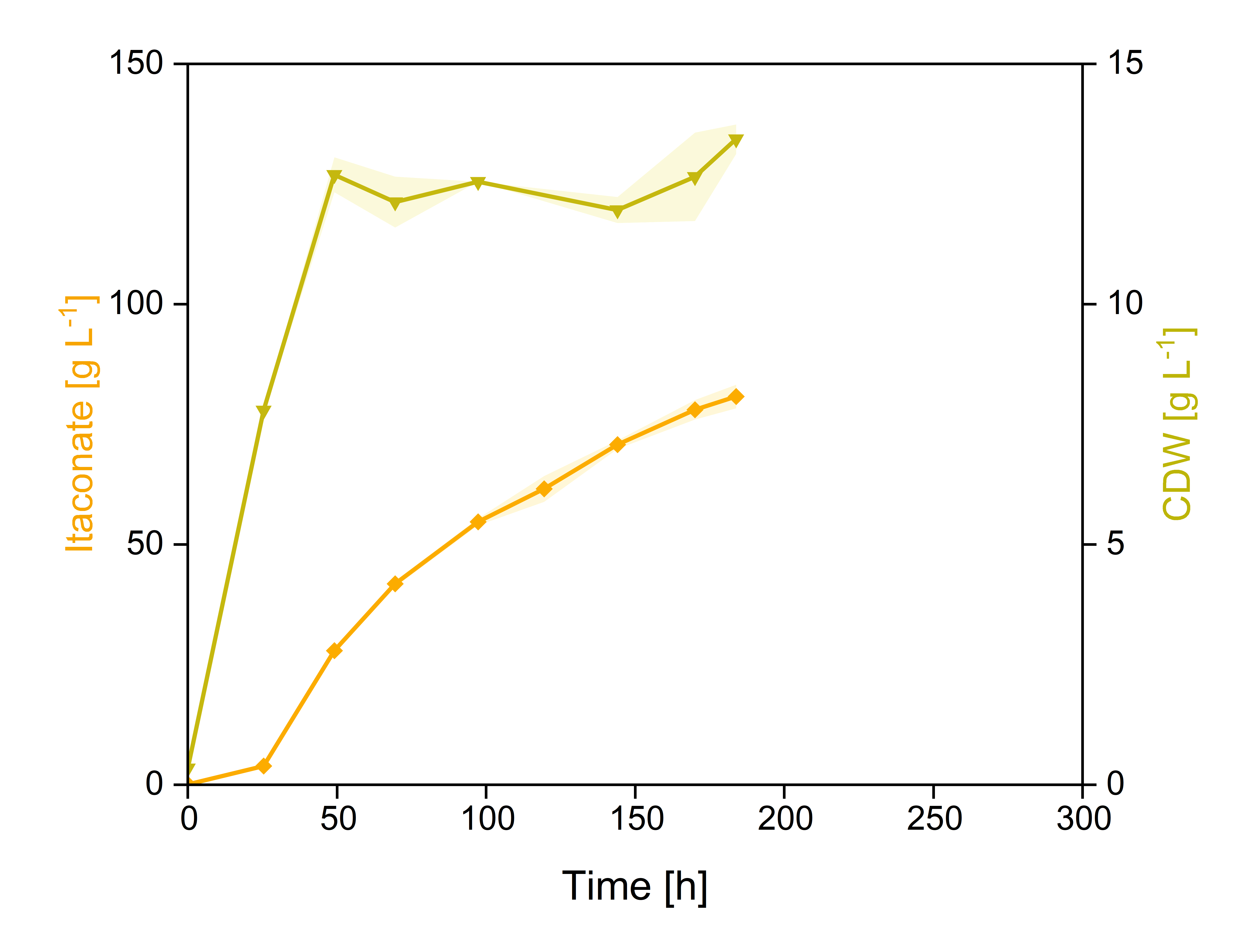

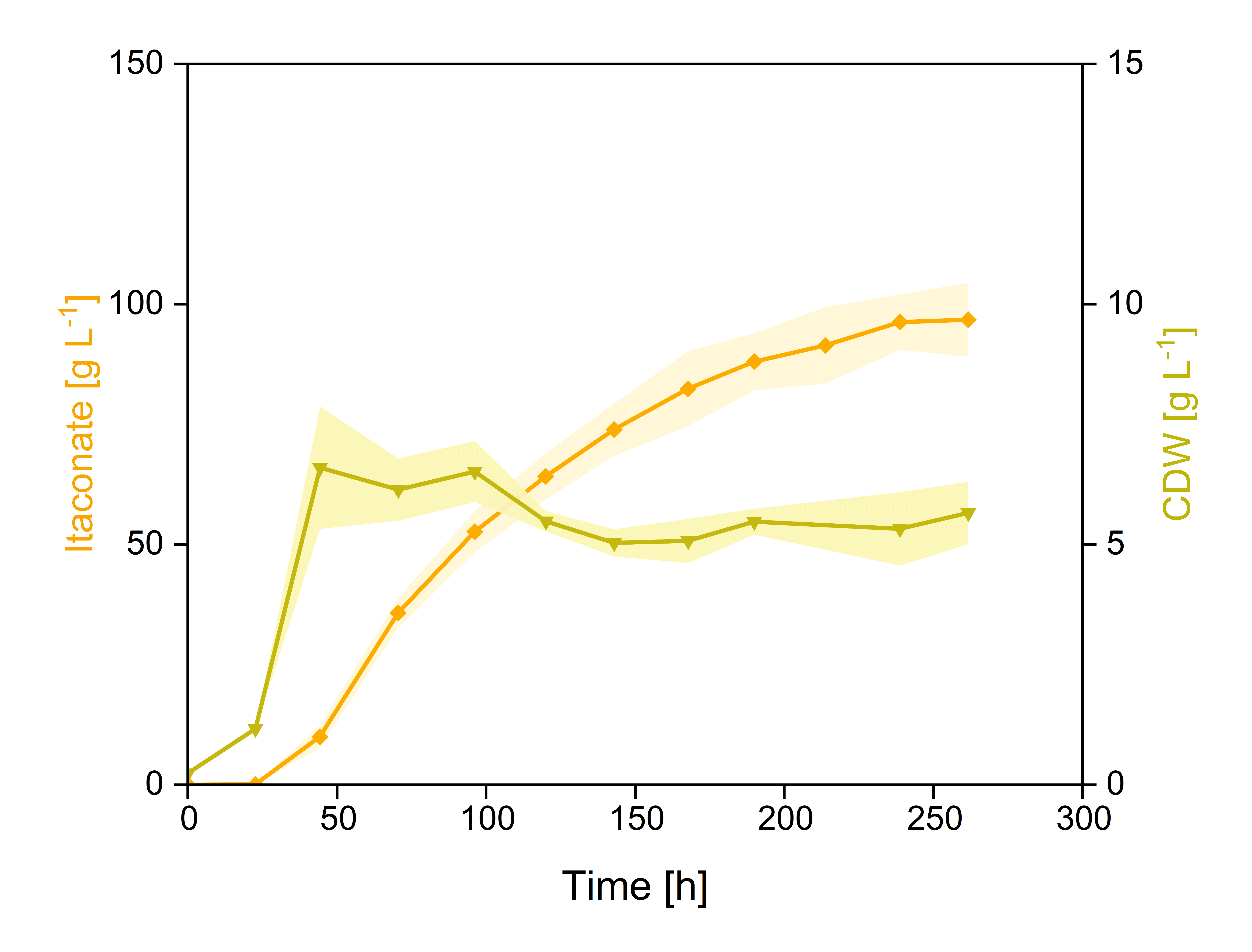
**Supplemental Figure 1.** Itaconate titers and CDW concentrations for the fed-batch bioreactor cultivation on acetate (A) and glucose (B). Itaconate concentration (orange squares), cell dry weight (CDW) (green triangles). Shades represent the minimum and maximum values of a biological duplicate (n=2).

**A**
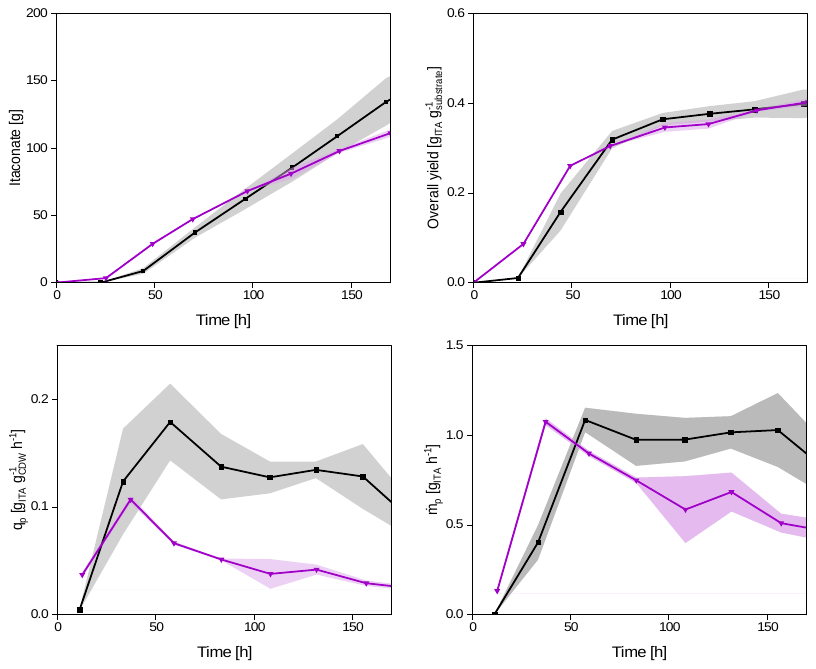

**BB**

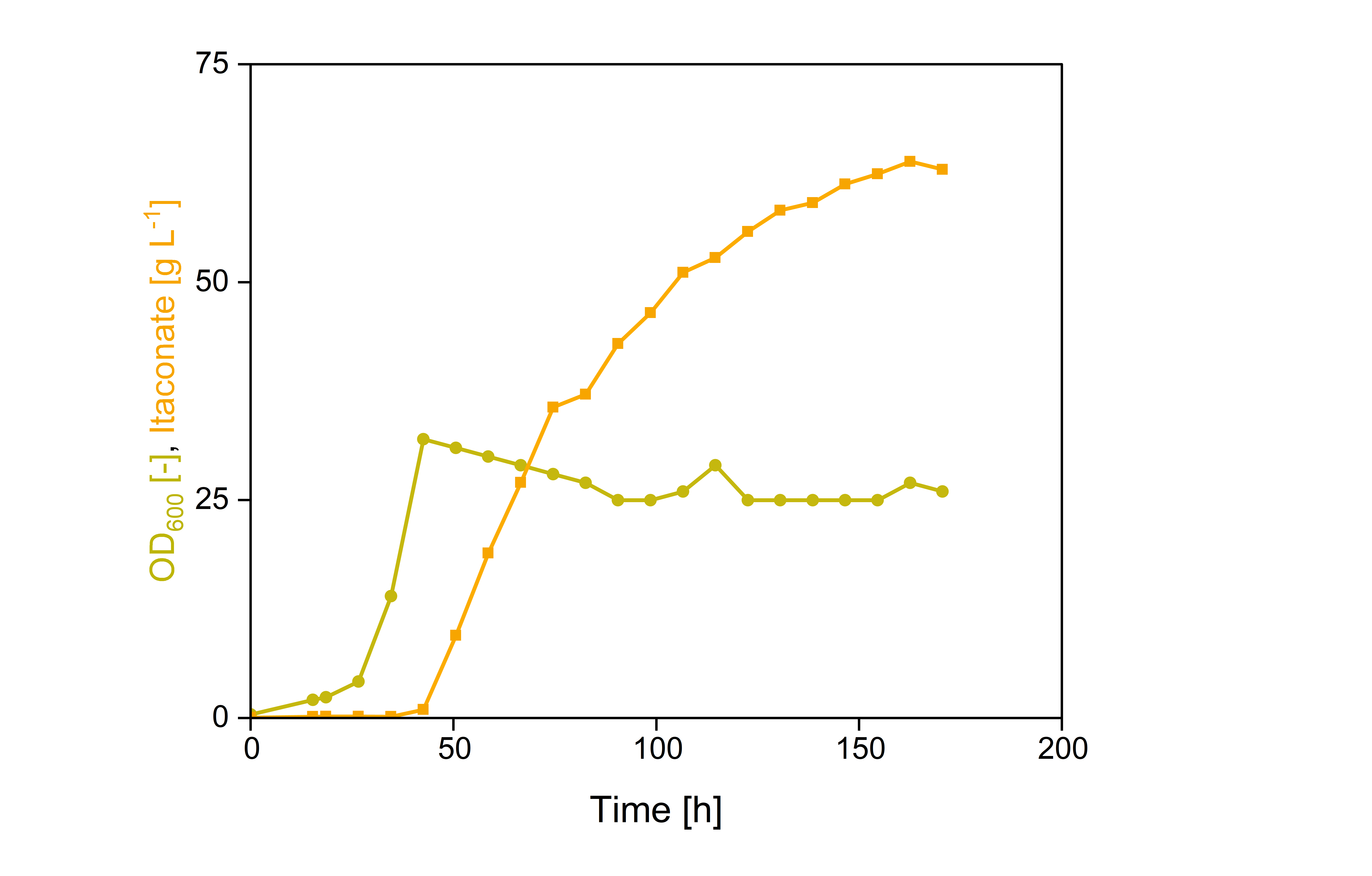

**Supplemental Figure 2.** Bioreactor cultivation of Ustilago maydis for itaconate production on acetic acid/potassium acetate using a pH and DO-coupled feeding strategy. Itaconate titer (orange squares) and optical density (OD) (green circles) are plotted over the cultivation time. The cultivation was performed at 30°C in MTM medium (50 mM MOPS, pH 6.8, 4 g L^-1^ NH_4_Cl) with dissolved oxygen levels between 70 and 80%. The pH was controlled with glacial acetic acid as titrant. DO-coupled fed shots with potassium acetate (500 g_ACE_ L^-1^), replenished the glucose concentration to 4 g L^-1^ in case of carbon depletion. Start volume of 500 mL, end volume of 518 mL. Itaconate production strain U. maydis ∆fuz7 ∆cyp3 ∆MEL ∆UA P_ria1_::P_etef_ P_etef_mttA was used as the production host. n =1.

**Supplementary Note S1: DO-coupled feeding script**

This script implements a DO-coupled feeding strategy. Feeding is triggered when the dissolved oxygen (DO) signal exceeds the defined threshold (DOTrg). A fixed feed shot is applied at 80% of the pump's maximum flow rate, followed by a delay phase until the next DO trigger. The script was executed in DASware Control. During cultivation, DOTrg was manually adjusted to approximately 6 DO units above the actual DO signal, and the feed-shot volume was increased over time to account for the increasing reactor volume.

```
Script for DO-coupled feeding strategy
DO-based feeding
Version 1.2, 2014/09/18, (c) DASGIP GmbH
Modified by Andreas Muesgens
Vessel Script

'Script parameters
Dim DOTrg As Double = 84               ' DO trigger [%]
Dim maxFeedFlowRate As Double = 200    ' Feed flow rate [mL/h]
Dim FeedShot As Double = 25            ' Feed shot volume [mL]
Dim DelayDuration_h As Double = 2      ' Delay between shots [h]

Dim Feed As Double = 0
Dim t_H  As Double
Dim t2_H As Double

If P IsNot Nothing Then
    With P
        Select Case .phase

            Case 0
                .phase = .phase + 1
                .LogMessage("feeding script active")

            Case 1
                If .DOPV > DOTrg Then
                    .phase = .phase + 1
                    .LogMessage("start DO based feeding event")
                End If

            Case 2
                Feed = maxFeedFlowRate
                t_H = .Runtime_h - .PhaseStart_H
                If t_H > (FeedShot / maxFeedFlowRate) Then
                    Feed = 0
                    .phase = .phase + 1
                    .LogMessage("feed DO based feeding stopped, waiting for DelayDuration")
                End If

            Case 3
                t2_H = .Runtime_h - .PhaseStart_H
                If t2_H > DelayDuration_h Then
                    .phase = .phase - 2
                    .LogMessage("Waiting for DOTrg")
                End If

        End Select

        .FCSP = Feed
    End With
End If
```
